# A novel risk score model based on seven protein-coding genes and three pseudogenes predicts overall survival in acute myeloid leukemia

**DOI:** 10.64898/2026.06.11.731554

**Authors:** Marco A. Fonseca-Montaño, Juan L. Ontiveros-Austria, José Alejandro Madrigal, Roberta Demichelis-Gómez, Laura Gómez-Romero, Iris K. Madera-Salcedo, Florencia Rosetti, José C. Crispín

**Affiliations:** Department of Immunology and Rheumatology, Instituto Nacional de Ciencias Médicas y Nutrición Salvador Zubirán, Mexico City, 14080, Mexico; Department of Hematology and Oncology, Instituto Nacional de Ciencias Médicas y Nutrición Salvador Zubirán, Mexico City, 14080, Mexico; Programa de Doctorado en Ciencias Biomédicas, Universidad Nacional Autónoma de México, Mexico City, 04510, Mexico; Escuela de Medicina y Ciencias de la Salud, Tecnologico de Monterrey, Monterrey, 64700, Mexico; Bioinformatics Department, Instituto Nacional de Medicina Genómica, Mexico City, 14610, Mexico

**Author notes:** Correspondence should be addressed to: José C. Crispín, MD, PhD, Vasco de Quiroga 15, Mexico City, 14080, Mexico.

**Keywords:** Acute myeloid leukemia, protein-coding genes, pseudogenes, overall survival, risk score model

## Abstract

Acute myeloid leukemia (AML) is the most common hematologic malignancy in adults and is associated with poor clinical outcomes. Accurate prognostic stratification remains essential for improving patient management and identifying potential therapeutic targets. Here, we analyzed RNA-seq data from the Therapeutically Applicable Research to Generate Effective Treatments (TARGET) and The Cancer Genome Atlas (TCGA) cohorts to identify genes associated with overall survival (OS) in patients with AML.

Using penalized regression modeling and integrative survival analyses, we identified a prognostic signature composed of seven protein-coding genes *(ACOT7, SLC35E4, SELPLG, CCND3, RRAS, ITGAX,* and *COMTD1*) and three processed pseudogenes (*FDPSP2, UBE2V1P13,* and *AL158214.1*) consistently associated with OS across independent AML cohorts. Based on these genes, we developed a novel risk score model that stratified AML patients into low- and high-risk groups with significantly different survival outcomes. The model demonstrated reproducible prognostic performance in TARGET and TCGA cohorts.

In addition, several genes included in the prognostic signature were significantly dysregulated in AML compared with healthy samples. Functional enrichment analyses further revealed that high-risk AML was associated with suppression of translational and RNA-processing pathways, together with cohort-specific enrichment of metabolic and immune-related programs.

Overall, this study presents a novel prognostic risk score integrating protein-coding genes and pseudogenes that robustly predicts OS in AML, providing a framework for improved prognostic stratification and potential biological insights into AML progression.

## INTRODUCTION

Acute myeloid leukemia (AML) is a heterogeneous hematological malignant disease caused by altered differentiation and dysregulated proliferation of myeloid blasts, that accumulate in the bone marrow and affect hematopoiesis (1). AML is the most frequent acute leukemia in adults, with a median age at diagnosis of ∼60 years and is frequently classified based on genetic abnormalities (1–4). Notably, AML remains a significant health challenge with a 5-year overall survival (OS) rate below 30%, which reflects its biological complexity and resistance to therapy associated with diverse genomic dysregulations (1,2).

The human genome has three major gene categories: protein-coding genes, long non-coding RNAs (lncRNAs), and pseudogenes, which together account for 24%, 45%, and 18% of all reported human genes, respectively (5). Previous studies have associated genomic alterations and dysregulated gene expression patterns with cancer development and clinical outcomes. Transcriptional dysregulation in cancer is a consequence of diverse genetic alterations in *trans*-factors (transcription factors, chromatin remodelers, cofactors) and *cis* elements (enhancers, promoters, insulators), that establish aberrant transcriptional networks that sustain cancer identity, a phenomenon referred to as transcriptional addiction, in which tumor cells become dependent (6,7).

The implications of genomic and transcriptional dysregulations in AML and other types of cancer have become indispensable for identifying biomarkers that refine prognostic assessment and therapeutic response (1,6,7). In this regard, previous studies have demonstrated that alterations in certain protein-coding genes, including *ASXL1, CEBPA, DNMT3A, FLT3, GATA2, IDH1/2, KIT,* and *NPM1*, confer a significant prognostic impact in patients with AML (1,2). In this regard, gene expression signatures have emerged as robust predictors of clinical outcomes. Ng and colleagues reported a 17-gene stemness signature that strongly correlated with minimal residual disease, relapse, and OS in patients across AML independent cohorts (8). Furthermore, previous reports have revealed prognostic signatures for risk stratification of AML patients based on genes related to genomic instability, disulfidptosis, ferroptosis, and necroptosis (9–13).

Beyond protein-coding genes, dysregulated lncRNA genes and pseudogenes are gaining recognition as non-traditional biomarkers in AML. For instance, the overexpression of *HOTAIRM1* or *LINC01268* is associated with poor clinical outcomes, whereas *MEG3* and *NEAT1* are frequently silenced in high-risk AML (14–17). On the other hand, the expression of pseudogenes, such as *VIM2P, POU5F1B, BMI1P1,* and *DUSP5P1,* has been proposed to have prognostic significance in AML (18–21). In addition, Jian and colleagues reported a gene signature based on 7 pseudogenes that predicted survival in patients with AML (22). Together, these findings highlight the AML transcriptome as a multi-layered prognostic reservoir.

Although several genes have been implicated in AML prognosis, the clinical significance of many remains unknown. In this context, accurately predicting the outcomes of patients is critical for guiding therapeutic decisions and improving survival. This study aimed to identify prognostic markers associated with OS in AML cohorts from the Therapeutically Applicable Research to Generate Effective Treatments (TARGET) and The Cancer Genome Atlas (TCGA). We constructed a prognostic model using least absolute shrinkage and selection operator (LASSO) regression, a machine-learning approach, and validated its predictive performance in both AML cohorts. We present a novel prognostic model based on seven protein-coding genes and three pseudogenes that predict OS at 1, 2, and 3 years, suggesting potential clinical applicability in patients with AML.

## MATERIAL AND METHODS

### Data collection

RNAseq-derived gene expression profiles from patients with AML, as primary diagnosis, were retrieved from TARGET (n=769) and TCGA (n=84) projects via GDC Data Portal (https://portal.gdc.cancer.gov/). AML records with available clinical data, OS time, and OS status were included. Additionally, RNA-seq-derived gene expression profiles from 355 healthy samples in the TARGET-AML project were retrieved from the GDC Data Portal.

### Identification and prioritization of OS-associated prognostic genes

First, gene expression was normalized to log_2_(TPM+1), and only genes with expression variance > 0 were included. For each gene, patient-level expression was dichotomized into high- and low-expression groups based on the lower (0.25) and upper (0.75) quartiles for downstream analyses.

Univariate Cox regression analyses were performed across the major human gene categories to identify OS-associated prognostic genes in the TARGET-AML and TCGA-AML cohorts. Subsequently, exploratory multivariable Cox regression screening analyses were conducted within each gene category to reduce the number of prognostic candidates while preserving genes with consistent prognostic trends across cohorts. Genes shared between TARGET-AML and TCGA-AML cohorts and exhibiting concordant hazard ratio (HR) directionality were prioritized as candidate markers for prognostic model construction.

These analyses were performed using the *survminer* package (https://anaconda.org/channels/conda-forge/packages/r-survminer/overview). HR and corresponding 95% confidence intervals were calculated, and *p*<0.05 was considered statistically significant.

### Prognostic model construction and validation

The *glmnet* package (https://anaconda.org/channels/conda-forge/packages/glmnet/overview) was used to perform least absolute shrinkage and selection operator (LASSO)-penalized Cox regression analysis to reduce redundancy among correlated prognostic candidates and identify the most informative genes for prognostic model construction. We employed 10-fold cross-validation to minimize overfitting and determine the optimal penalization parameter (λ).

Genes exhibiting non-zero LASSO coefficients were retained for prognostic signature construction. For each patient, the risk score was calculated as a weighted linear combination of gene expression values and their corresponding LASSO coefficients according to the following formula:

*Risk score* = (LASSO coefficient of selected gene 1 × expression of selected gene 1) + … + (LASSO coefficient of selected gene *n* × expression of selected gene *n*).

Time-dependent receiver operating characteristic (ROC) analyses were performed using the *timeROC* package (https://anaconda.org/channels/conda-forge/packages/r-timeroc/overview) to calculate the area under the curve (AUC) at 1, 2, and 3 years of OS. LASSO coefficient profiles and cross-validation plots (partial likelihood deviance versus log(λ)) were also generated. TARGET-AML and TCGA-AML cohorts were used as training and validation cohorts, respectively.

### Survival analyses and evaluation of risk score differences

The median risk score was used as a cutoff to stratify AML patients into low- and high-risk groups. OS differences between groups were evaluated using Kaplan-Meier survival curves and the log-rank test. Tables showing the number of patients at risk were included. Statistical significance was defined as *p*<0.05. These analyses were performed using the *survival* and *survminer* packages.

Additionally, Mann-Whitney U tests were performed to identify statistically significant differences in the risk score between healthy and AML samples in the TARGET cohort. This test was also used to evaluate differences in risk score distributions between sexes and between independent AML cohorts.

### Cox proportional hazards regression analyses

Clinicopathological covariates, including diagnosis age, sex, mutation count, and risk score, were evaluated through univariate and multivariable Cox proportional hazards regression analyses to assess their associations with OS in the TARGET-AML and TCGA-AML cohorts. These analyses were performed using the *survival* and *survminer* packages. HRs with corresponding 95% confidence intervals were calculated, and *p*<0.05 was considered statistically significant. Forest plots were generated using the *forestplot* package (https://anaconda.org/channels/conda-forge/packages/r-forestplot/overview).

### Guilt-by-association analysis and functional annotation

For guilt-by-association analysis, we used log_2_(TPM+1) normalized gene expression from the TARGET-AML cohort to compute two-sided Pearson correlations between each pseudogene selected by LASSO analysis and the expression of all protein-coding genes. Then, we selected protein-coding genes that showed *r*≥0.5 and *p*<0.05 to perform Gene Ontology (GO) overrepresentation analysis for biological processes and molecular functions. These analyses were performed using the *clusterProfiler* package (https://anaconda.org/channels/bioconda/packages/bioconductor-clusterprofiler/overview). Statistically significant (*p*<0.05) GO biological processes and molecular functions terms were shown in dot plots.

### Gene set enrichment analyses

The *DESeq2* (https://anaconda.org/channels/bioconda/packages/bioconductor-deseq2/overview) and *clusterProfiler* packages were used to perform gene set enrichment analysis (GSEA), independently, in the TARGET-AML and TCGA-AML cohorts using a pre-ranked strategy to identify biological pathways associated with risk stratification. Genes were ranked based on continuous statistics reflecting both the magnitude and direction of differential expression between high- and low-risk AML groups, enabling the detection of coordinated transcriptional programs across the entire expression spectrum. Positive and negative ranking values indicated enrichment toward high- and low-risk groups, respectively. GSEA was conducted using GO, and statistical significance was assessed following correction for multiple testing. Enrichment results were summarized using normalized enrichment scores (NES) and adjusted *p*-values, and were primarily visualized using dot plots, where gene ratio, statistical significance, and gene set size were simultaneously represented to facilitate comparative interpretation between high- and low-risk groups.

### Spearman correlation analyses

Gene expression levels of 16 clinically relevant AML-associated genes (*FLT3, IDH1, IDH2, NPM1, CEBPA, DDX41, TP53, ASXL1, BCOR, EZH2, RUNX1, SF3B1, SRSF2, STAG2, U2AF1,* and *ZRSR2*) were compared between low- and high-risk AML groups in the TARGET and TCGA cohorts using the Mann–Whitney U test. Subsequently, Spearman correlation analyses were performed to assess the association between gene expression levels and the risk score in both cohorts. Correlation coefficients (*R*) and corresponding *p*-values were calculated for each gene across samples. *p*<0.05 was considered statistically significant.

### Ethical statement

The data used in this study were obtained from publicly available databases. Since these databases consist of anonymized data that is freely accessible to the public, no ethical approval or informed consent was required for this study. The data were used solely for research purposes and were in compliance with the relevant ethical guidelines for the use of publicly available data.

## RESULTS

### Construction and validation of a cross-cohort prognostic risk score in AML

To identify OS-associated prognostic markers in AML patients, we performed univariate Cox regression analyses across three major gene categories: protein-coding genes, lncRNAs, and pseudogenes. In the TARGET-AML cohort, 519 protein-coding genes, 188 lncRNAs, and 892 pseudogenes were significantly associated with a protective effect (HR<1, *p*<0.05), whereas 3281 protein-coding genes, 1298 lncRNAs, and 619 pseudogenes were associated with increased risk (HR>1, *p*<0.05; Fig 1A–C). Similarly, in the TCGA-AML cohort, 895 protein-coding genes, 656 lncRNAs, and 650 pseudogenes showed protective associations, while 1438 protein-coding genes, 281 lncRNAs, and 156 pseudogenes exhibited risk-associated effects (Fig 1D–F).

**Fig 1.**
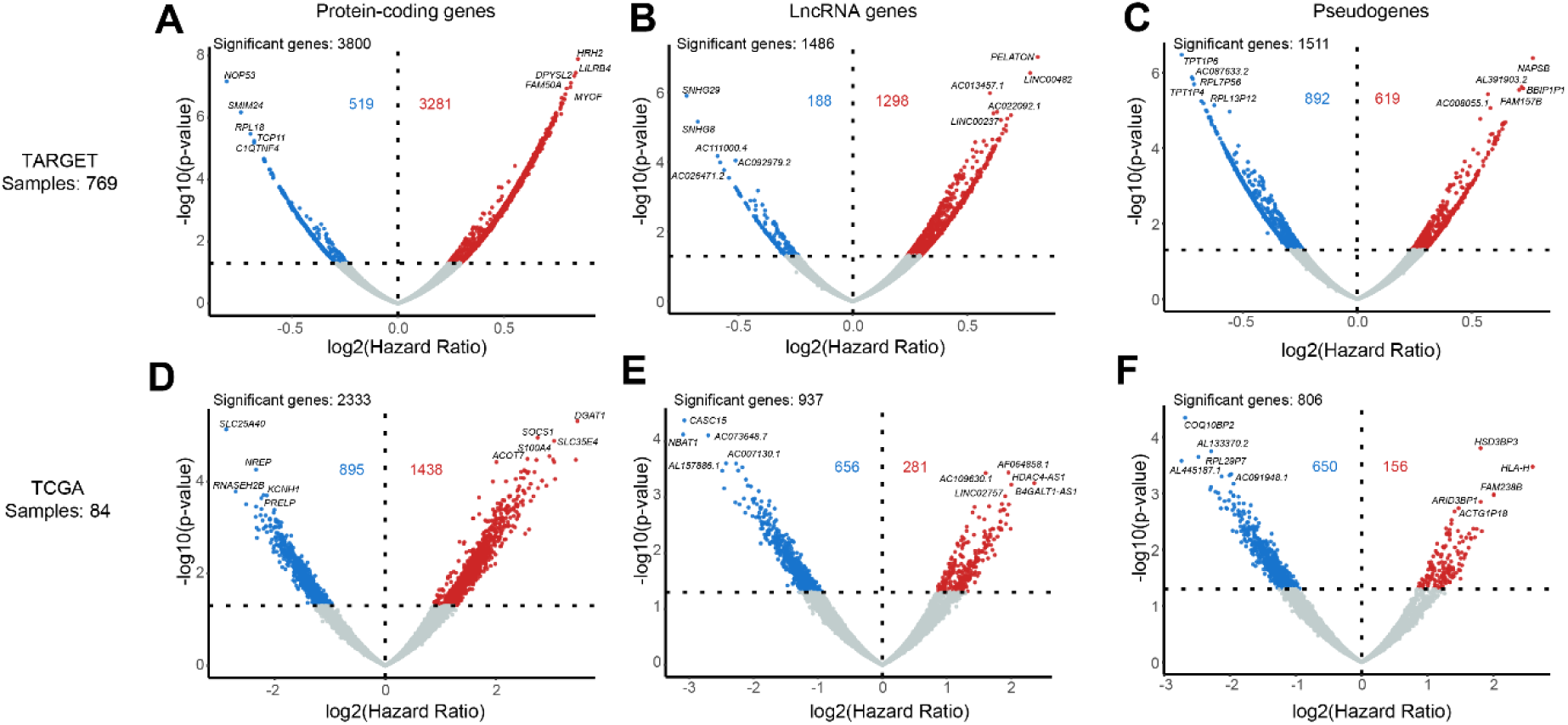
Univariate Cox regression identifies OS–associated genes across gene categories in independent AML cohorts. Volcano plots of genes associated with OS in the TARGET cohort (n = 769) for (A) protein-coding genes, (B) lncRNA genes, and (C) pseudogenes, and in the TCGA cohort (n = 84) for (D) protein-coding genes, (E) lncRNA genes, and (F) pseudogenes. The x-axis represents log_2_(HR), and the y-axis shows −log10(*p*-value). Horizontal and vertical dashed lines indicate the significance threshold (*p*<0.05) and log_2_(HR) = 0, respectively. Genes with HR<1 are shown in blue (protective association), whereas genes with HR>1 are shown in red (risk association).

To further prioritize robust OS-associated candidates across independent AML cohorts, we subsequently performed an exploratory multivariable Cox regression screening analysis within each gene category. This approach reduced the number of candidate prognostic markers while preserving genes with consistent prognostic trends between TARGET and TCGA cohorts (Fig 2A–B). Cross-cohort intersection analyses identified 19 protein-coding genes, 8 lncRNAs, and 8 pseudogenes shared between both cohorts (Fig 2C–E). A final set of 14 OS-associated candidate genes exhibiting concordant HR directionality across cohorts was subsequently prioritized for penalized regression modeling.

**Fig 2.**
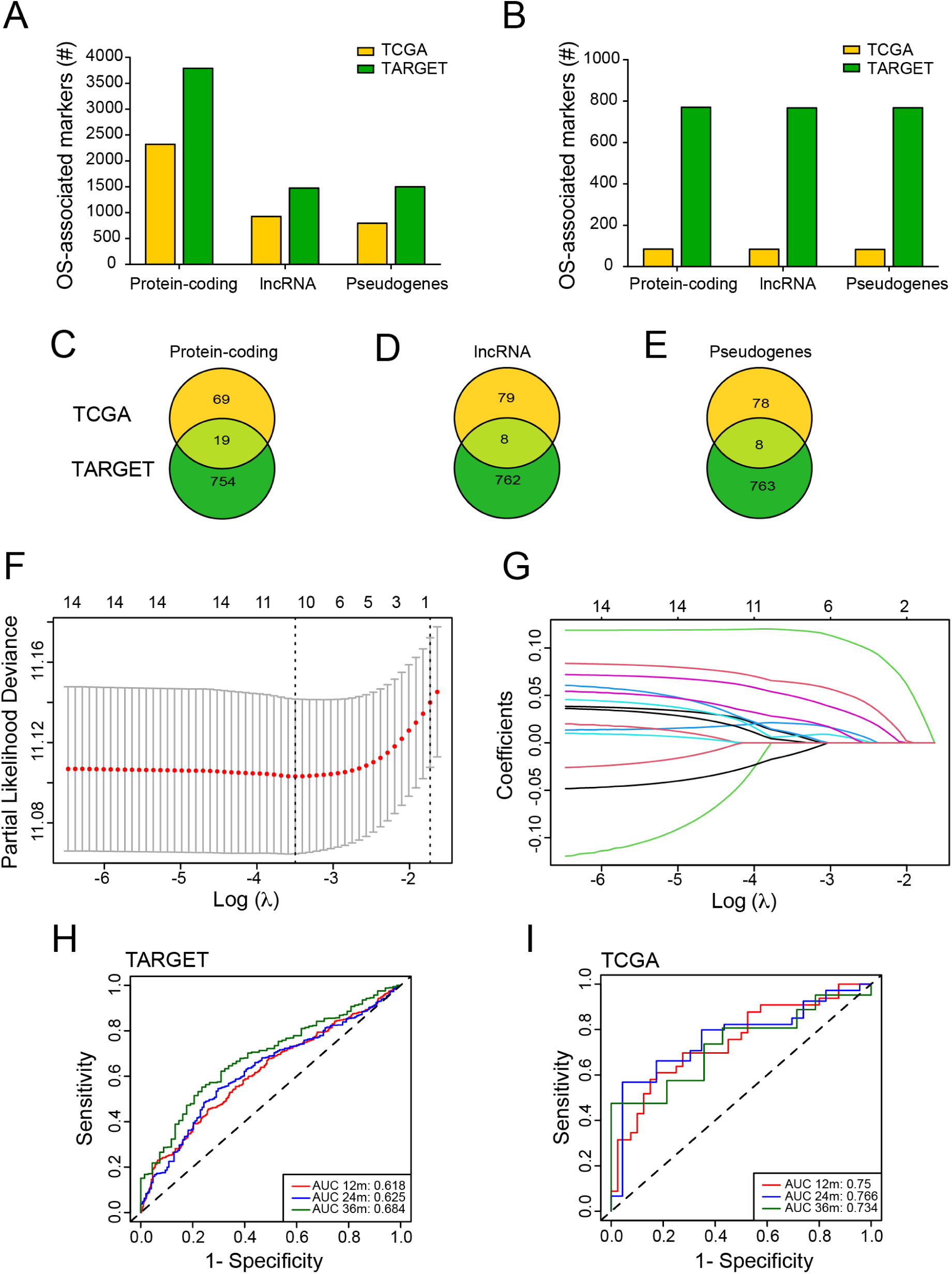
Cross-cohort prioritization and LASSO-based construction of the AML prognostic risk score. (A) Number of genes significantly associated with OS identified by univariate Cox regression across gene categories in the TARGET and TCGA cohorts. (B) Number of candidate prognostic genes prioritized through exploratory multivariable Cox regression screening analyses across gene categories in both cohorts. Venn diagrams showing the overlap of OS-associated prognostic markers between TARGET and TCGA cohorts for (C) protein-coding genes, (D) lncRNA genes, and (E) pseudogenes. (F) Cross-validation plot showing partial likelihood deviance as a function of log(λ) in the TARGET cohort. Dashed vertical lines indicate the optimal penalization parameters. (G) LASSO coefficient profiles of prognostic candidate genes across log(λ) values in the TARGET cohort. Time-dependent ROC curves evaluating the prognostic performance of the risk score model at 1, 2, and 3 years in the (H) TARGET and (I) TCGA cohorts. AUC values are indicated for each time point.

Given the potential redundancy and collinearity among prognostic candidate genes, LASSO-Cox regression analysis was employed to reduce model complexity and identify the most informative predictors for prognostic model construction. According to the partial likelihood deviance and lambda optimization analyses, the optimal prognostic model consisted of 10 genes (Fig 2F–G), including seven protein-coding genes (*ACOT7, SLC35E4, SELPLG, CCND3, RRAS, ITGAX,* and *COMTD1*) and three pseudogenes (*UBE2V1P13, FDPSP2,* and *AL158214.1*). The resulting prognostic signature generated the following risk score model:

*Risk score* = [(0.00841280055356348) x *ACOT7* expression] + [(0.0635529008109001) x *SLC35E4* expression] + [(0.0204739530089515) x *SELPLG* expression] + [(0.00708508770012495) x *CCND3* expression] + [(0.0494564151416561) x *RRAS* expression] + [(0.00191092854764465) x *ITGAX* expression] + [(0.118872597312218) x *COMTD1* expression] + [(0.0101944746419609) x *UBE2V1P13* expression] + [(0.0251928645078268) x *FDPSP2* expression] + [(−0.0121142448180268) x *AL158214.1* expression].

To evaluate the prognostic performance of the resulting risk score, time-dependent ROC analyses were performed in both cohorts. In the TARGET-AML cohort, the model achieved AUC values of 0.618, 0.625, and 0.684 at 1, 2, and 3 years, respectively (Fig 2H). Independent validation in the TCGA-AML cohort yielded AUC values of 0.750, 0.766, and 0.734 at the same time points (Fig 2I). Collectively, these results support the robustness, reproducibility, and cross-cohort prognostic performance of the resulting AML risk signature.

### Prognostic value for clinicopathological covariates and risk score in AML

We performed univariate Cox regression analyses for demographic and clinicopathological covariates (diagnosis age, sex, and mutation count) together with the risk score. We observed that the risk score emerged as a significant prognostic factor for OS in both the TARGET-AML and TCGA-AML cohorts (Fig 3A–B). Notably, the prognostic significance of the risk score persisted after multivariable Cox regression adjustment in both cohorts.

**Fig 3.**
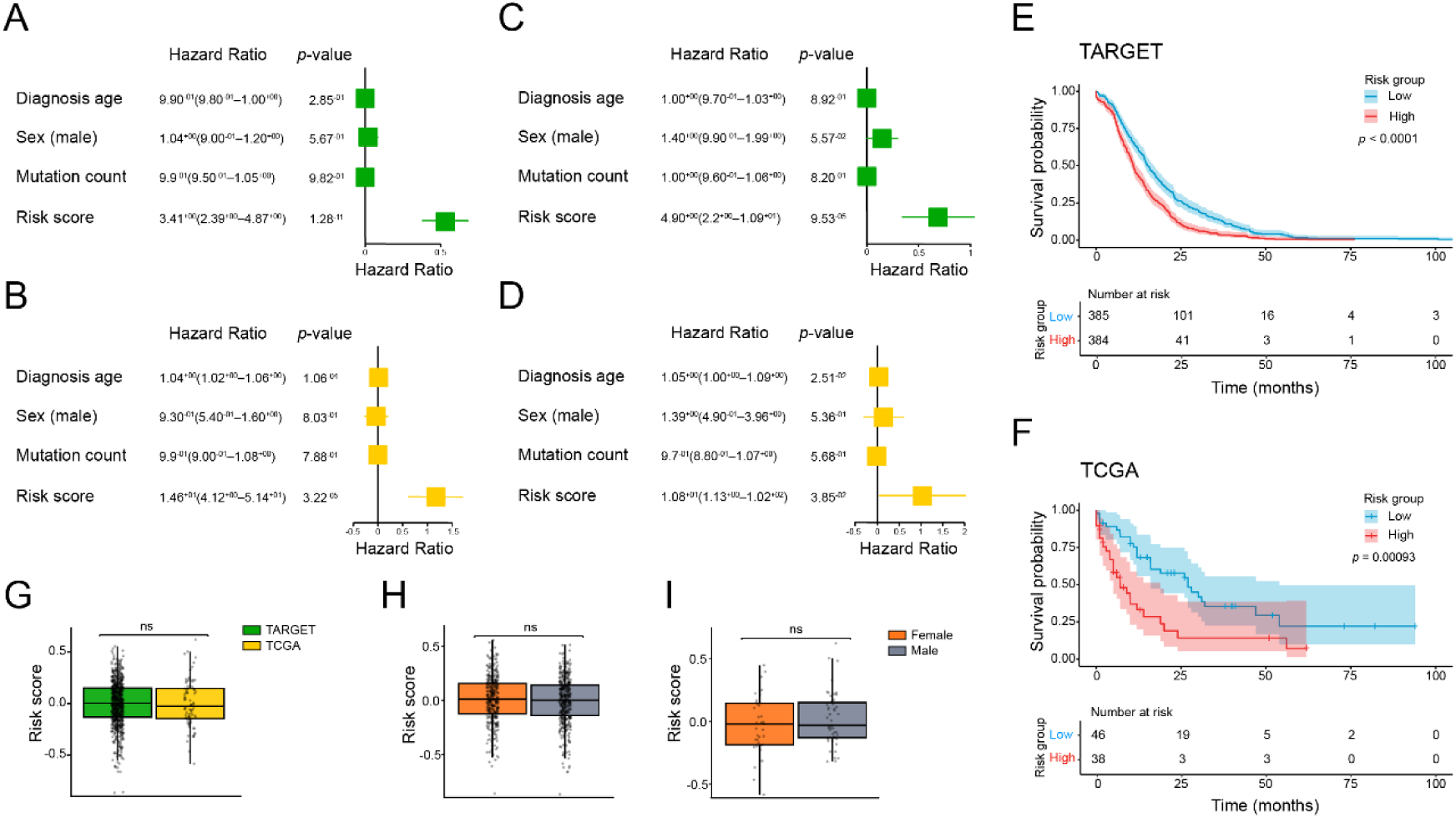
Prognostic value of clinicopathological variables and the risk score in independent AML cohorts. Forest plots from univariate Cox regression analyses of clinicopathological variables and risk score in the (A) TARGET and (B) TCGA cohorts. Forest plots from multivariate Cox regression analyses in the (C) TARGET and (D) TCGA cohorts. HR with corresponding *p*-values are shown. Kaplan–Meier survival curves comparing OS between low- and high-risk groups in the (E) TARGET and (F) TCGA cohorts. Shaded areas represent confidence intervals, and numbers at risk are indicated below each plot. (G) Distribution of risk scores between TARGET and TCGA cohorts. (H–I) Distribution of risk scores according to sex in the (H) TARGET and (I) TCGA cohorts. Statistical significance is indicated as follows: **** (*p*<1 × 10⁻⁴), *** (*p*<1 × 10⁻³), ** (*p*<1 × 10⁻²), * (*p*<0.05), and ns (*p*≥0.05).

Furthermore, age at diagnosis also behaved as an independent prognostic factor for OS in the TCGA-AML cohort (Fig 3C–D).

AML patients from the TARGET-AML and TCGA-AML cohorts were subsequently stratified into low- and high-risk groups according to the median risk score. Kaplan-Meier survival analyses demonstrated that low-risk AML patients exhibited significantly better OS compared with high-risk patients in the TARGET-AML cohort. These findings were consistently validated in the TCGA-AML cohort (Fig 3E–F).

Notably, no statistically significant differences in the risk score distribution were observed between TARGET-AML and TCGA-AML cohorts (Fig 3G). Likewise, no significant sex-associated differences in the risk score were detected in either cohort (Fig 3H–I). Collectively, these findings further highlight the relevance and consistency of the risk score across independent cohorts.

### Functional annotation for pseudogenes that compose the risk score

The function of the 7 protein-coding genes included in the risk score (*ACOT7*, *SLC35E4, SELPLG, CCND3, RRAS, ITGAX,* and *COMTD1*) has been reported. On the other hand, previous studies have demonstrated that pseudogenes play essential roles in cancer biology and could serve as potential prognostic markers. However, their functional and clinical roles have not been widely explored as protein-coding and lncRNA genes. In this context, we aimed to bioinformatically characterize the molecular functions and biological processes associated with the expression of the 3 pseudogenes included in the risk score (*FDPSP2, UBE2V1P13,* and *AL158214.1*) using guilt-by-association analysis. For *UBE2V1P13,* we identified a significant correlation with the expression of 1428 protein-coding genes, and subsequent GO over-representation analysis revealed enrichment for diverse molecular functions, including histone-modifying activity, histone binding, ATP-dependent activity acting on DNA, helicase activity, structural constituent of chromatin, and DNA helicase activity (Fig 4A). For *FDPSP2,* we detected a significant correlation with the expression of 1,662 protein-coding genes, and subsequent GO over-representation analysis revealed enrichment for molecular functions, including ubiquitin-like protein transferase activity, ubiquitin-protein transferase activity, ubiquitin protein ligase activity, and protein serine kinase activity (Fig 4B). In contrast, we found a significant correlation between *AL158214.1* and the expression of 6 protein-coding genes, and subsequent GO overrepresentation analysis revealed enrichment for molecular functions, including growth factor receptor binding, growth factor activity, and glycosaminoglycan binding (Fig 4C). Further analysis showed that protein-coding genes correlated with *UBE2V1P13* are enriched in biological processes, including nuclear chromosome segregation, DNA replication, mitotic nuclear division, and DNA-templated DNA replication (Fig 4D). For protein-coding genes correlated with *FDPSP2,* the enriched biological processes include Golgi vesicle transport, endosomal transport, vacuolar transport, and peptidyl-serine modification (Fig 4E). Finally, the protein-coding genes correlated with *AL158214.1* showed enrichment for biological processes, including cell chemotaxis to fibroblast growth factor, regulation of cell chemotaxis to fibroblast growth factor, regulation of endothelial cell chemotaxis, and endothelial cell chemotaxis (Fig 4F). These results suggest the potential roles for the 3 pseudogenes in the AML context; however, further functional studies are required to validate this hypothesis.

**Fig 4.**
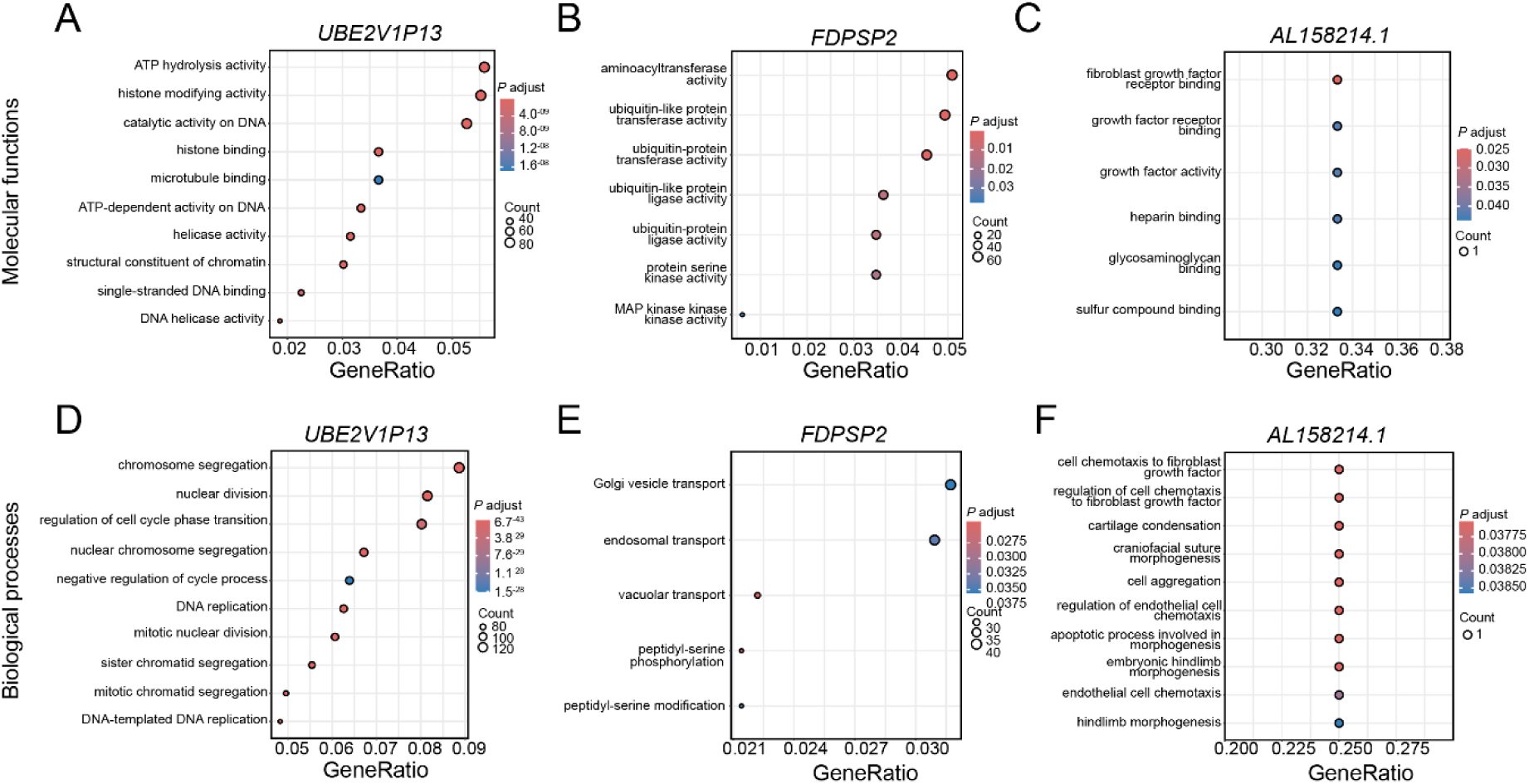
GO over-representation analysis of protein-coding genes associated with pseudogenes included in the risk score. Dot plots showing enriched GO terms for protein-coding genes correlated with the expression of *UBE2V1P13, FDPSP2,* and *AL158214.1*. (A–C) Molecular function and (D–F) biological process categories. The x-axis represents GeneRatio, dot size indicates gene count, and color denotes adjusted *p*-value.

### Risk score shows differences between healthy and AML samples

Although the risk score was developed to stratify risk groups based on OS in AML, we explored its behavior in healthy (n=355) and AML samples (n=769) available in the TARGET cohort. Notably, the expression of *SLC35E4, SELPLG, CCND3, ITGAX, COMTD1, FDPSP2, UBE2V1P13,* and *AL158214.1* was significantly decreased in AML patients when compared to healthy samples. Interestingly, *ACOT7* and *RRAS* expression did not show significant differences between healthy and AML samples (Fig 5A). This expression pattern was corroborated by finding that AML patients have lower risk scores than healthy samples from the TARGET cohort (Fig 5B). These results suggest global transcriptional differences between healthy and AML conditions, explaining the behavior previously stated.

**Fig 5.**
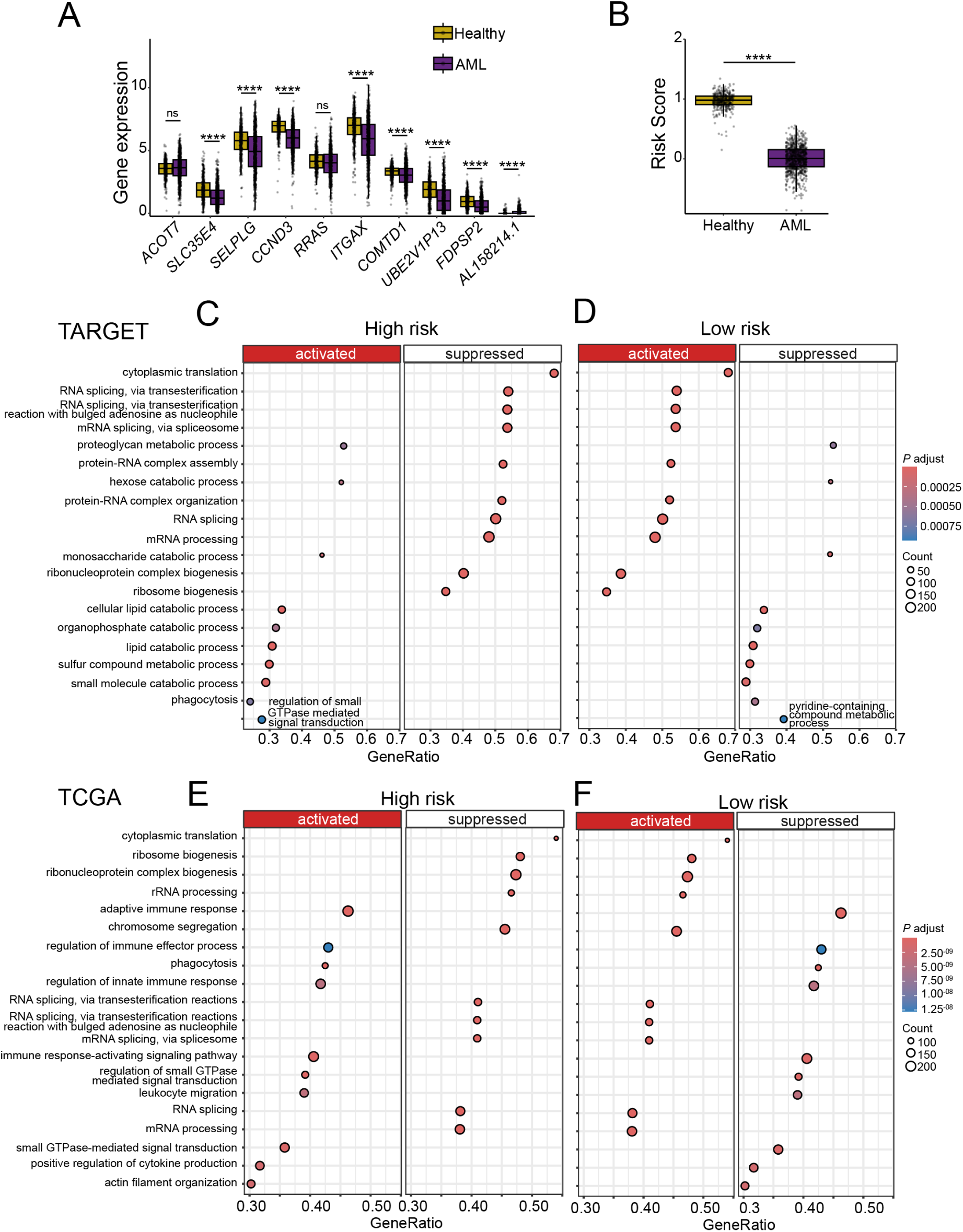
Expression differences, risk score distribution, and GSEA across AML cohorts. (A) Gene expression levels of selected prognostic markers in healthy and AML samples from the TARGET cohort. (B) Distribution of the risk score between healthy and AML samples from the TARGET cohort. GSEA results comparing (C) high- and (D) low-risk groups in the TARGET cohort. GSEA results comparing (E) high- and (F) low-risk groups in the TCGA cohort. Dot plots display enriched pathways separated into positively enriched (activated) and negatively enriched (suppressed) categories. The x-axis represents GeneRatio, dot size indicates gene count, and color denotes adjusted *p*-value. Statistical significance is indicated as follows: **** (*p* < 1 × 10⁻⁴), *** (*p*<1 × 10⁻³), ** (*p*<1 × 10⁻²), * (*p*<0.05), and ns (*p*≥0.05).

### Suppression of translational and RNA-processing programs defines high-risk AML

GSEA revealed consistent pathway programs associated with risk stratification across the TARGET-AML and TCGA-AML cohorts. In this context, a dominant feature of the high-risk AML in both cohorts was the suppression of translational and RNA-processing pathways, including cytoplasmic translation, ribosome biogenesis, ribonucleoprotein complex biogenesis, RNA splicing, and mRNA processing (Fig 5C and E). Particularly, in the TARGET cohort, this suppression co-occurred with positive enrichment of catabolic and metabolic processes (small molecule, lipid, organophosphate, and monosaccharide catabolism) (Fig 5D). In the TCGA cohort, high-risk AML instead showed enrichment of immune- and signaling-related pathways, including adaptive and innate immune responses, leukocyte migration, cytokine production, phagocytosis, and small GTPase–mediated signaling, together with cytoskeletal organization (Fig 5F). Conversely, the low-risk AML in both cohorts displayed the reciprocal pattern, with enrichment of translational and RNA-processing programs and relative suppression of catabolic pathways in TARGET and immune/migratory programs in TCGA cohort. Overall, the convergence across cohorts indicates that loss of ribosomal and RNA-processing activity is a hallmark of high-risk AML, while low-risk AML retains these core programs. This suggests that high-risk AML is characterized by a shift away from coordinated gene expression and protein synthesis toward alternative states, including metabolic adaptation (TARGET) or immune and migratory signaling (TCGA), which may be essential in AML biology of this risk group.

### Elevated expression of clinically actionable AML genes associates with high-risk disease

Finally, we assessed whether the expression of 16 genes commonly used for mutation screening, diagnosis, and therapeutic stratification in AML differed between low- and high-risk AML patients. Across TARGET and TCGA cohorts, we observed that *IDH1, DDX41, ASXL1,* and *SF3B1* were consistently and significantly upregulated in high-risk patients, while showing lower expression levels in low-risk groups (Fig 6A and B).

**Fig 6.**
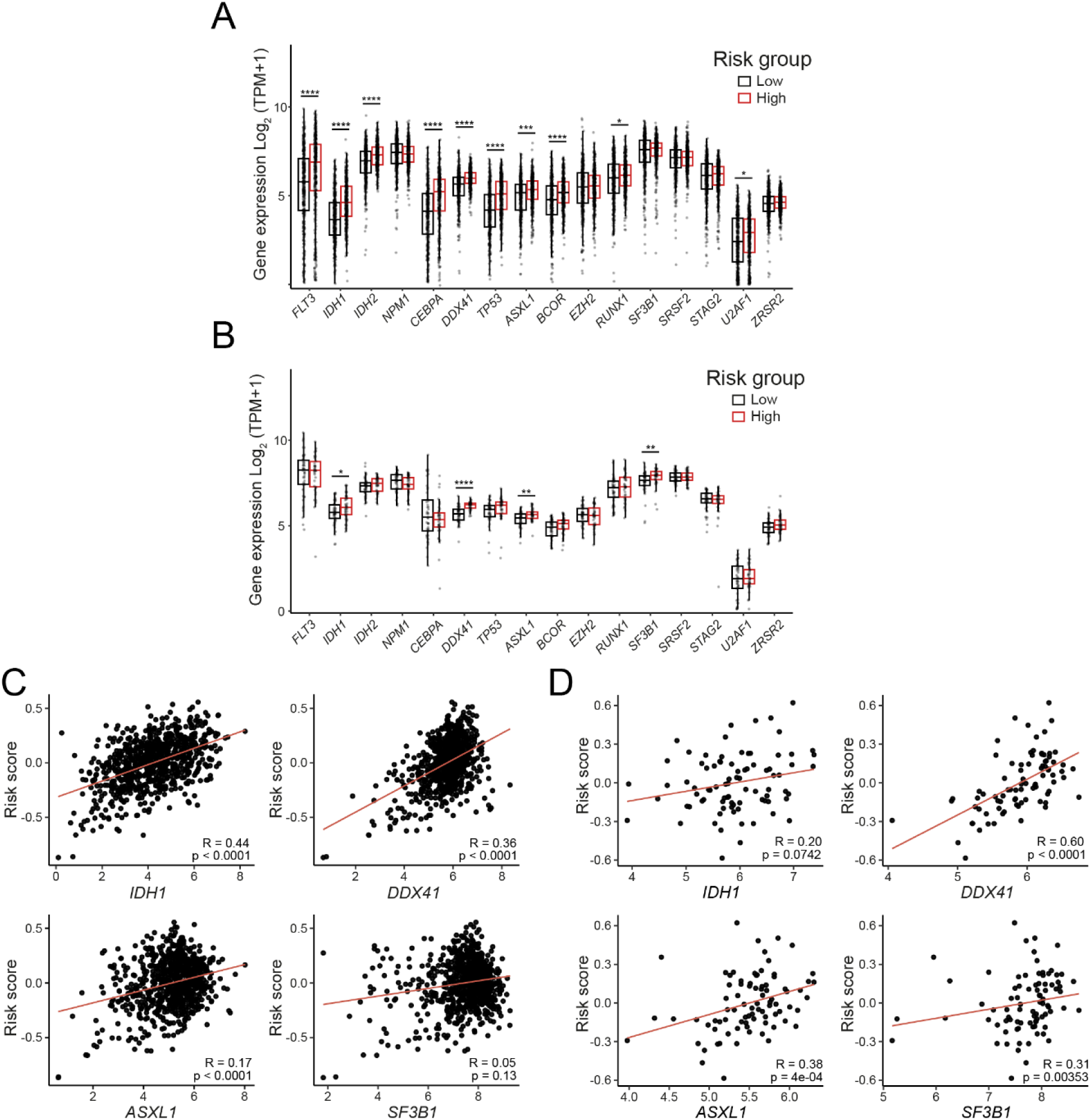
Expression and correlation of clinically relevant AML-associated genes with the risk score. Gene expression levels of 16 clinically relevant AML-associated genes in low-and high-risk AML groups in the (A) TARGET and (B) TCGA cohorts. Expression values are shown as log₂(TPM+1). Spearman correlation between gene expression and the risk score in the (C) TARGET and (D) TCGA cohorts. Each point represents an individual sample. Correlation coefficients (R) and corresponding *p*-values are indicated. Statistical significance is denoted as **** (*p*<1 × 10⁻⁴), *** (*p*<1 × 10⁻³), ** (*p*<1 × 10⁻²), * (*p*<0.05), and ns (*p*≥0.05).

To further evaluate the relationship between gene expression and risk stratification, we performed correlation analyses between gene expression levels and the risk score. Notably, *DDX41* and *ASXL1* exhibited a robust and consistent positive correlation with the risk score across both cohorts, whereas *IDH1* and *SF3B1* showed more variable or cohort-dependent associations (Fig 6C and D). Collectively, these findings indicate that increased expression of selected AML-associated genes, particularly *DDX41* and *ASXL1*, is closely linked to high-risk disease, supporting their potential relevance as molecular features associated with adverse prognosis.

## DISCUSSION

AML constitutes the most common type of acute leukemia in adults and is associated with poor clinical outcomes. Despite progress in diagnosis and treatment, the survival rate for AML remains remarkably low, and accurate prognostic evaluation is essential to improving OS in patients. In this regard, AML research has focused on identifying gene markers that predict clinical outcomes or serve as potential therapeutic targets (1–4). In our study, by performing univariate and multivariate regression analysis, we identified seven protein-coding genes (*ACOT7, SLC35E4, SELPLG, CCND3, RRAS, ITGAX,* and *COMTD1*) and three processed pseudogenes (*FDPSP2, UBE2V1P13,* and *AL158214.1*) with a consistent prognostic role associated with OS in AML patients across independent cohorts. Then, using LASSO regression analysis, we developed a novel prognostic risk score model that includes the 10 genes mentioned above and categorizes AML patients into low- and high-risk groups, with better and worse OS, respectively. Notably, the expression of *SLC35E4, SELPLG, CCND3, ITGAX, COMTD1, FDPSP2, UBE2V1P13,* and *AL158214.1* was significantly increased in AML compared with healthy samples, suggesting that their downregulation may have a relevant role in AML biology.

Among the genes mentioned above, *ACOT7* belongs to the Acyl-CoA thioesterase family, which catalyzes the hydrolysis of fatty acyl-CoAs. In cancer, *ACOT7* depletion induces cytostasis in the absence of a DNA damage response, promotes cell cycle arrest, and activates the PKCζ–p53–p21 signaling pathway *in vitro*. Moreover, its prognostic role has been evaluated in breast and lung cancer, demonstrating that low *ACOT7* expression is associated with improved OS (23). In contrast, high *ACOT7* expression is associated with reduced event-free survival and OS in AML patients (24). *SLC35E4* is a putative transporter, and previous studies have suggested that it may be involved in ceRNA networks in cholangiocarcinoma (25,26); however, its role in AML biology remains elusive.

*SELPLG* is an SLe(x)-type proteoglycan that interacts with E-, P-, and L-selectins to mediate rolling of leukocytes over vascular surfaces during the early stage of inflammation. Previous reports have shown that the prognostic value of *SELPLG* in OS, disease-specific survival, disease-free, and progression-free intervals varies across cancer types (27). Cheng and colleagues assessed the prognostic role of diverse cell adhesion molecule genes in AML and found that *SELPLG* expression negatively correlates with OS in AML patients (28). *CCND3* is a D-type cyclin that plays a key role in the cell cycle by controlling progression from G1 to S phase, and its deregulation and prognostic significance have been reported in solid cancers (29). Recurrent mutations in *CCND3* confer resistance to apoptosis, decreased cell-cycle arrest, and increased proliferation in the presence of FLT3 inhibitors in AML (30). Furthermore, Matsuo and colleagues reported increased *CCND3* expression and mutations in the PEST domain, which stabilizes cyclin D3 and promotes cell proliferation in *MLL*-rearranged AML (31). In addition, a recent study showed that *CCND3* expression is a prognostic marker in AML patients and is upregulated in relapsed/refractory disease (32). *RRAS* belongs to the Ras family of GTPases, and previous reports have demonstrated that *RRAS* plays oncogenic roles (33–36). However, its prognostic role across cancer types remains unclear. *ITGAX* belongs to the integrin family, which is essential for cell adhesion, migration, and immune surveillance, and its deregulation has been linked to cancer promotion (37,38). In this context, *ITGAX* has been identified as a cancer gene hub associated with metastasis and with prognostic significance across diverse types of solid cancers (39,40). Also, *ITGAX* has been reported to promote gastric cancer progression through the epithelial-mesenchymal transition pathway (37). Wang and colleagues found that *ITGAX* is an angiogenic promoter, inducing high VEGF-A expression and activating VEGFR2 signaling via the PI3K/Akt-mediated c-Myc pathway *in vitro* and *in vivo* (41). Notably, previous studies in AML showed that *ITGAX* expression correlates with CTLA4 expression (42), and high *ITGAX* expression was associated with reduced OS (43). On the other hand, *COMTD1* is a putative O-methyltransferase, and previous studies have developed prognostic models that include *COMTD1* for breast cancer and esophageal squamous cell carcinoma (44,45). However, the precise functional role of *COMTD1* remains unknown across different types of cancer, including AML.

Several evidence has exposed the relevance of processed pseudogenes in cancer biology, and their expression has recently been explored in the clinical context (14,15,17). However, few studies about pseudogenes have been reported in AML. In this regard, a previous study showed that *VDAC1P8* pseudogene is upregulated in AML, and 24 transcription factor binding sites (MYC, GATA1/3, FL11, IRF4, RUNX1) were identified in the target regulatory region of *VDAC1P8*, some of which had been associated with leukemogenesis (46). *TPTEP1* was identified as a potential tumor suppressor due to its *in vitro* role in controlling AML cell growth by inactivating the JNK/c-JUN signaling pathway (47). A recent study found that *TPTEP1* regulates the malignant behavior of AML cells by interacting with miR-4295 and upregulating GADD45α, a target of miR-4245 that exerts a suppressive effect on AML cells (48). On the other hand, low *VIMP2P* and *BMI1P1*, and high *DUSP5P1* expression have been associated with poor clinical outcome in AML patients (18,49,50). In contrast, the high *POU5F1B* expression is associated with improved OS in AML patients (51). Notably, the expression of *VDAC1P8, DUSP5P1, VIM2P, BMI1P1, POU5F1B,* and *TPTEP1* differs between AML and healthy controls (46–51). In our research, we report, for the first time, that the *UBE2V1P13, FDPSP2,* and *AL158214.1* pseudogenes are mainly associated with DNA-, ubiquitin-, and growth-factor-related processes, respectively, suggesting their potential relevance in the AML context. However, further functional experimental assays may be essential to validate their role *in vitro* or *in vivo*.

Notably, the transcriptional and pathway-level features captured by the risk score are consistent with previously reported mechanisms underlying cancer progression. In particular, the suppression of translational and RNA-processing programs observed in high-risk AML aligns with growing evidence that alterations in RNA splicing, ribosome biogenesis, and mRNA processing are critical drivers of tumor development and progression. Dysregulation of these processes has been widely associated with diverse hallmarks in multiple cancer types, including hematological malignancies (52–55). In this context, the consistent enrichment of translational and RNA-processing pathways in low-risk patients may reflect a more preserved transcriptional state, whereas their suppression in high-risk AML suggests a shift toward alternative functional programs. Notably, these patterns were accompanied by cohort-specific features, including metabolic pathway enrichment in TARGET and immune- and signaling-related processes in TCGA, supporting the notion that high-risk AML is characterized by distinct but convergent biological features.

This study has certain limitations. First, the prognostic risk score model primarily relies on retrospective RNA-seq bulk data from the TARGET and TCGA cohorts. In this regard, prospective transcriptomic data from AML patients across additional independent cohorts may help determine whether the prognostic value of our risk score model is limited to specific cohorts in which ancestry and ethnicity might influence results or confirm its broader applicability. In addition, both datasets lack comprehensive clinical information regarding several recurrent genetic alterations commonly used for AML prognostic stratification. Therefore, future studies incorporating these molecular features may help determine whether specific genetic alterations are associated with our risk score model. Second, reliance on bioinformatic analysis may necessitate *in vitro* or *in vivo* experiments to accurately validate and understand the functional roles of genes in our risk score model, some of which remain unclear in AML biology, such as *SLC35E4, COMTD1, FDPSP2, UBE2V1P13,* and *AL158214.1*. Finally, although the model shows significant prognostic stratification at 1, 2, and 3 years of OS, its clinical usefulness requires validation through prospective trials that evaluate its ability to inform treatment decisions. Despite these considerations, our predictive model offers promising insights toward improved diagnosis and the identification of potential new therapeutic targets in AML patients.

## ACKNOWLEDGMENTS

This work was funded by a grant from Fundación Gonzalo Río Arronte (S794) to Alejandro Madrigal. This work was partially performed at cluster INMEGEN and received technical support from Walter Santos. The authors would like to thank Dr. Adrián Soto for his advice.

## AUTHOR CONTRIBUTIONS

**Conceptualization:** Marco A. Fonseca-Montaño, Alejandro Madrigal, Roberta Demichelis-Gómez, José C. Crispín

**Investigation:** Marco A. Fonseca-Montaño, Laura Gómez-Romero.

**Methodology:** Marco A. Fonseca-Montaño, Laura Gómez-Romero, José C. Crispín.

**Data curation:** Marco A. Fonseca-Montaño.

**Formal analysis:** Marco A. Fonseca-Montaño.

**Validation:** Marco A. Fonseca-Montaño, Laura Gómez-Romero.

**Visualization:** Marco A. Fonseca-Montaño, Iris K. Madera-Salcedo, Florencia Rosetti.

**Supervision:** Laura Gómez-Romero, Florencia Rosetti, Roberta Demichelis-Gómez, José C. Crispín.

**Writing – original draft:** Marco A. Fonseca-Montaño.

**Writing – review & editing:** Marco A. Fonseca-Montaño, Juan L. Ontiveros-Austria, José Alejandro Madrigal, Roberta Demichelis-Gómez, Laura Gómez-Romero, Iris K. Madera-Salcedo, Florencia Rosetti, José C. Crispín.

## Notes

### Competing Interest Statement

The authors have declared no competing interest.

